# The α-tubulin of *Laodelphax striatellus* facilitates the passage of rice stripe virus (RSV) and enhances horizontal transmission

**DOI:** 10.1101/502831

**Authors:** Yao Li, Danyu Chen, Jia Hu, Lu Zhang, Yin Xiang, Fang Liu

**Affiliations:** College of Horticulture and Plant Protection, Yangzhou University, Yangzhou, China; State Key Laboratory for Biology of Plant Diseases and Insect Pests, Institute of Plant Protection, Chinese Academy of Agricultural Sciences, Beijing, China; Joint International Research Laboratory of Agriculture & Agri-Product Safety, Yangzhou University, Yangzhou, China; Jiangsu Co-Innovation Center for Modern Production Technology of Grain Crops, Yangzhou University, Yangzhou, China

## Abstract

Rice stripe virus (RSV), causal agent of rice stripe disease, is transmitted by the small brown planthopper (SBPH, *Laodelphax striatellus*) in a persistent manner. The midgut and salivary glands of SBPH are the first and last barriers in viral circulation and transmission, respectively; however, the precise mechanisms used by RSV to cross these organs and re-inoculate rice have not been fully elucidated. We obtained full-length cDNA of *L. striatellus α-tubulin 2* (*LsTUB*) and found that RSV infection increased the level of LsTUB in vivo. Furthermore, LsTUB was shown to bind the RSV nonstructural protein 3 (NS3) in vitro. RNAi was used to reduce *LsTUB* expression, which caused a significant reduction in RSV titer, *NS3* expression, RSV inoculation rates, and transmission to healthy plants. Electrical penetration graphs (EPG) showed that LsTUB knockdown by RNAi did not impact SBPH feeding; therefore, the reduction in RSV inoculation rate was likely caused by the decrease in RSV transmission. These findings suggest that LsTUB mediates the passage of RSV through midgut and salivary glands and leads to successful horizontal transmission.

## Introduction

The survival of plant viruses is largely dependent on the efficient transmission to plant hosts by viral-specific vector(s) [1,2]. Insects transmit over 70% of all known plant viruses [3]. Hemipteran insects (e.g. leafhoppers, planthoppers, aphids and whiteflies), transmit approximately 55% of insect-vectored plant viruses; as these insects have distinctive piercing-sucking mouthparts with needle-like stylet bundles that are comprised of two maxillary and two mandibular stylets [3–5]. Four categories of insect vector - plant virus transmission relationships have been described as follows: non-persistent; semi-persistent; persistent: propagative and persistent: circulative [2,6]. Plant viruses with persistent relationships enter vectors via the gut, pass through various tissues and ultimately reach the salivary glands and ovaries. Viruses are transmitted horizontally from the salivary glands of the vector into healthy plants and are vertically transmitted from female ovaries to offspring [7]. Barriers to the persistent transmission of plant viruses in insect vectors include the following: (i) midgut infection barriers; (ii) dissemination barriers, including midgut escape and salivary gland infection barriers; (iii) salivary gland escape barriers; and (iv) transovarial transmission barriers [8,9]. A deeper understanding of the mechanistic basis of virus transmission through these four barriers will facilitate the development of novel methods to control the systemic spread of plant viruses [10].

Previous studies demonstrated that the persistent transmission of viruses in different insect tissues requires specialized interactions between components of the virus and vector. For example, in the aphid *Myzus persicae*, the coat protein read-through domain (CP-RTD) of *Beet western yellows virus* binds Rack-1 and membrane-bound glyceraldehyde-3-phosphate dehydrogenase; these processes are thought to facilitate transcytosis of luteoviruses in the aphid midgut and accessory salivary glands [11]. The coat proteins of *Tomato leaf curl New Delhi virus* and *Cotton leaf curl Rajasthan virus* were shown to interact with *Bemisia tabaci* midgut protein to facilitate trafficking of viral particles from the midgut into the insect hemolymph [12]. Furthermore, the *Rice ragged stunt virus* nonstructural protein Pns10 interacted with the *Nilaparvata lugens* oligomycin-sensitivity conferral protein to enhance virus titer in salivary gland cells [13]. Such interactions in different insect vectors are highly complex and diverse, and their effect on the horizontal transmission of viruses remains unclear.

Rice stripe virus (RSV, genus *Tenuivirus*) has inflicted severe yield losses in rice throughout East Asia [14,15]. RSV is transmitted by the small brown planthopper (SBPH), *Laodelphax striatellus*, in a persistent, circulative-propagative manner. Once inside SBPH, RSV invades the midgut epithelium to establish infection and then spreads into various SBPH tissues through the hemolymph. Recently, molecular interactions between RSV and various SBPH tissues have received increased attention [16]. In the midgut, a direct interaction between the nonstructural protein NS4 and the nucleocapsid protein (CP) of RSV promoted viral spread in viruliferous SBPH [17]. Furthermore, the interaction between the SBPH sugar transporter 6 (LsST6) and RSV CP was shown to be essential for RSV transfer across the midgut infection barrier [18]. The interaction between RSV CP and SBPH vitellogenin (LsVg) facilitated vertical transmission of the virus [19]. Further work revealed that *LsVg* expression has tissue-specificity, and that LsVg produced in hemocytes was responsible for vertical transmission of RSV [20]. We previously reported that RSV was horizontally transmitted to rice plants via salivation during the feeding of insect vectors [21]. A series of salivary-specific transcriptome and proteome analyses revealed numerous genes involved in RSV transmission [22,23]; however, only cuticular protein (CPR1) and a G protein pathway suppressor 2 (GPS2) impacted RSV transmission and replication in salivary glands [23,24]. Obviously, the mechanism that RSV uses to overcome the salivary gland barrier and then undergo horizontal transmission to the plant warrants further investigation.

In this study, proteomics was used to show that α-tubulin was highly expressed in viruliferous SBPH as compared with naïve SBPH, which suggests that *L. striatellus* tubulin (LsTUB) may have a role in mediating RSV transmission. We show that LsTUB facilitated the passage of RSV through the midgut and salivary gland barriers and enhanced viral transmission from SBPH to rice. Yeast two-hybrid and pull-down assays provided evidence that the interaction of LsTUB and the RSV nonstructural protein NS3 likely constitutes a critical step in RSV transmission.

## Results

### cDNA cloning and sequence analysis of *LsTUB*

Based on proteomic analysis of SBPH salivary glands, 33 differentially expressed proteins were identified in viruliferous and virus-free, naïve SBPH (data not shown). The *L. striatellus* Tubulin α-2 (LsTUB) was among the 33 differentially expressed proteins; this is significant partly because tubulin heterodimers are known to function in virus assembly and transport [25–27]. Using the conserved sequence of *Tubulin α-2* from NCBI (GenBank accession no. AY550922.1) as an *in silico* probe, full-length cDNA sequences of *LsTUB* (1658 bp, GenBank accession no. KF934411) were identified and cloned from female SBPH adults. *LsTUB* contained a 1353-bp ORF that encoded a putative protein of 450 amino acids, a 94-bp 5’ untranslated region (UTR), and a 212-bp 3’ UTR. The translated cDNA of *LsTUB* yields a protein with a mass of approximately 50.0 kDa, and theoretical isoelectric point (pI) of 5.01. SMART analysis showed that LsTUB contains two conserved domains, including a GTPase domain (amino acids 49-246) with a GDP-binding site (amino acids 142-147) and a C-terminal domain (amino acids 248-393) (Fig 1A). Alignment of the LsTUB predicted protein sequence with other TUB proteins indicated a high level of identity with Hemipteran TUB proteins, including NlTUB in *Nilaparvata lugens* (GenBank accession no. ANJ04673.1, 100% identity) and LlTUB in *Lygus lineolaris* (GenBank accession no. AHG54247.1, 99% identity) (Fig. 1B).

**Figure 1.**
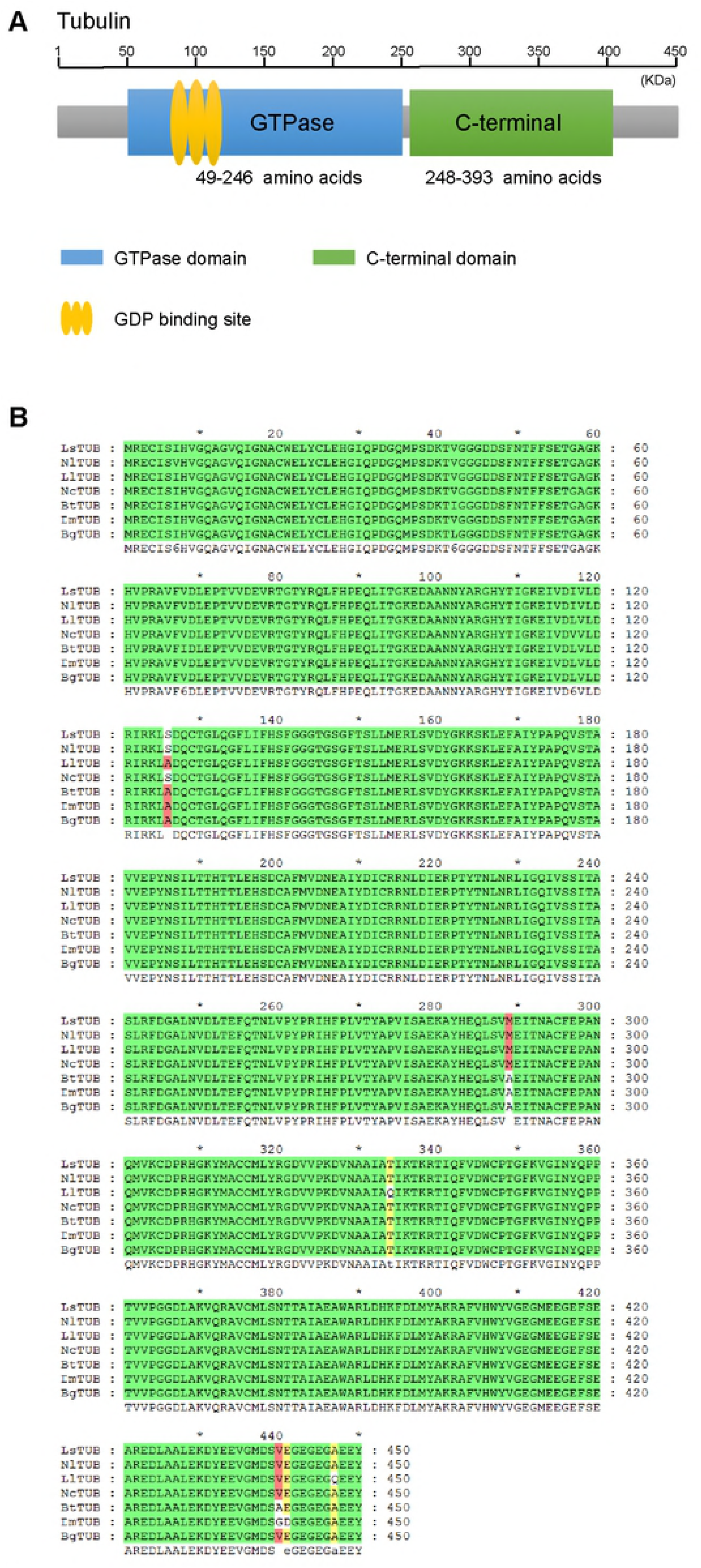
LsTUB protein structure and amino acid alignment. (A) Exons (gray rectangle) and conserved domains in LsTUB are shown, including GTPase and C-terminal domains in blue and green rectangles, respectively. The GDP-binding site of LsTUB is marked with three golden ellipses. (B) Deduced amino acid sequence alignments of TUB in seven insect species; alignments were constructed using ClustalW software. Green shading indicates conserved tubulin residues in seven insect species; red and yellow shading indicates species-specific residues. Abbreviations indicate tubulin from the following insect species: LsTUB, *L. striatellus*; NlTUB, *Nilaparvata lugens*; LlTUB, *Lygus lineolaris*; NcTUB, *Nephotettix cincticeps*; BtTUB, *Bombus terrestris*; DmTUB, *Drosophila melanogaster*; and BgTUB, *Blattella germanica*.

### RSV infection increases *LsTUB* expression

To further evaluate differential expression of *LsTUB* in viruliferous vs naïve SBPH, qRT-PCR and Western blot analysis were conducted to quantify mRNA and protein expression levels, respectively. The mRNA expression levels of *LsTUB* were significantly up-regulated in viruliferous SBPH (Fig. 2A). The trend in gene expression was consistent with changes in protein expression as determined by immunoblotting with anti-LsTUB antisera (Fig. 2B).

**Figure 2.**
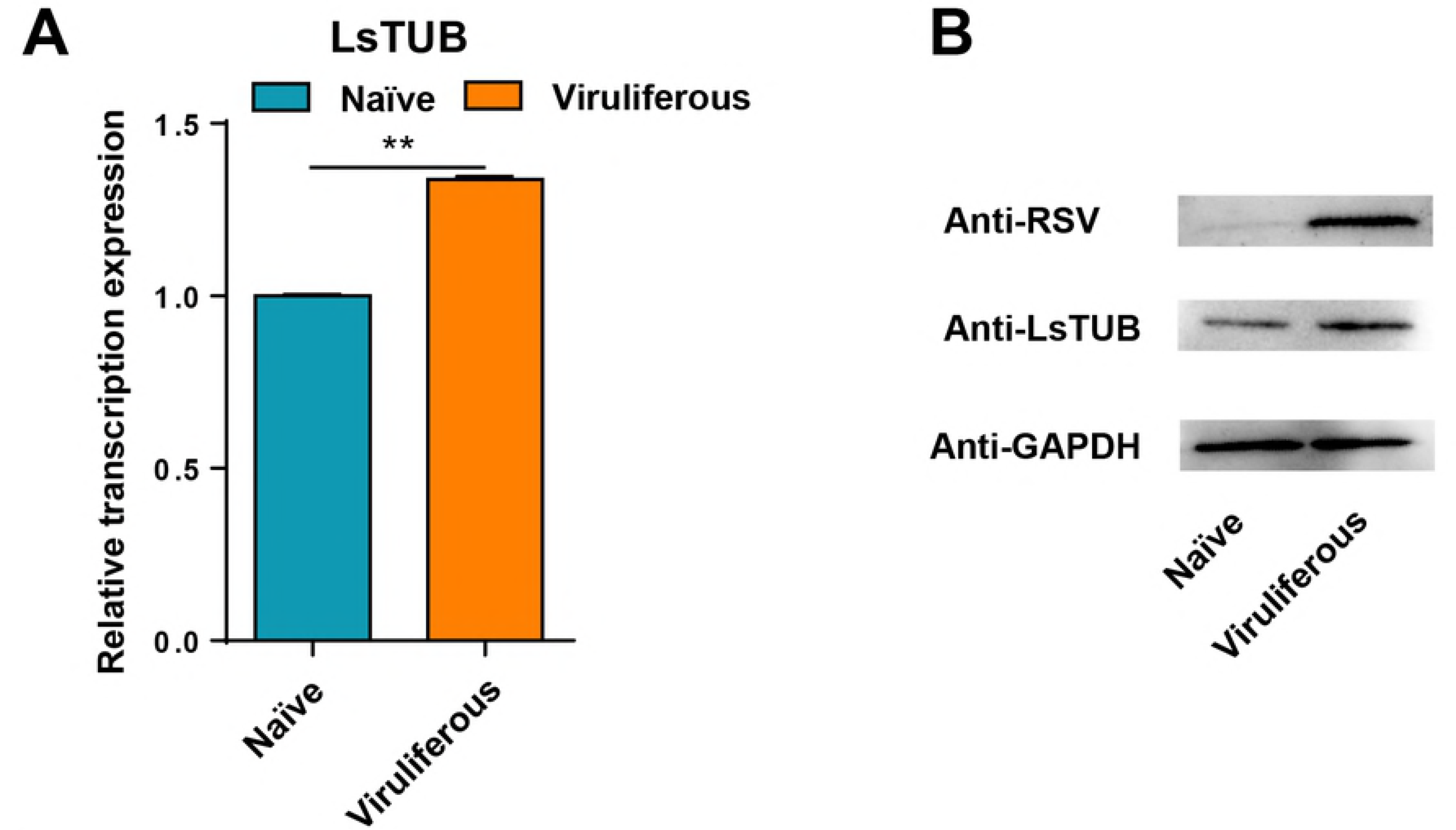
mRNA and protein expression levels of *LsTUB* in naïve and viruliferous SBPH. (A) qRT-PCR analysis of *LsTUB* expression in naïve and RSV-infected SBPH. Treatments were replicated three times. Means ± S.E. T-test analysis: \**P*<0.05, \*\**P*<0.01, and \*\*\**P*<0.001. (B) Western blot analysis of LsTUB production in naïve and viruliferous SBPH. GAPDH was used as control.

### LsTUB co-localizes with RSV in different tissues

Our results suggest that *LsTUB* is expressed at higher levels in response to RSV in viruliferous SBPH; thus, we investigated whether this response coincides with RSV transmission. In this experiment, monoclonal antisera of LsTUB and RSV CP were labeled with Alexa Fluor 555 and 488, respectively. In SBPH midgut (Fig. 3A-A’’’), hemocytes (Fig. 3B-B’’’) and principal salivary glands (Fig. 3C-C’’’), LsTUB and RSV CP co-localized (see arrows, Fig 3A’-D’’’’).The results indicated that LsTUB and RSV accumulate and co-localize throughout the SBPH body.

**Figure 3.**
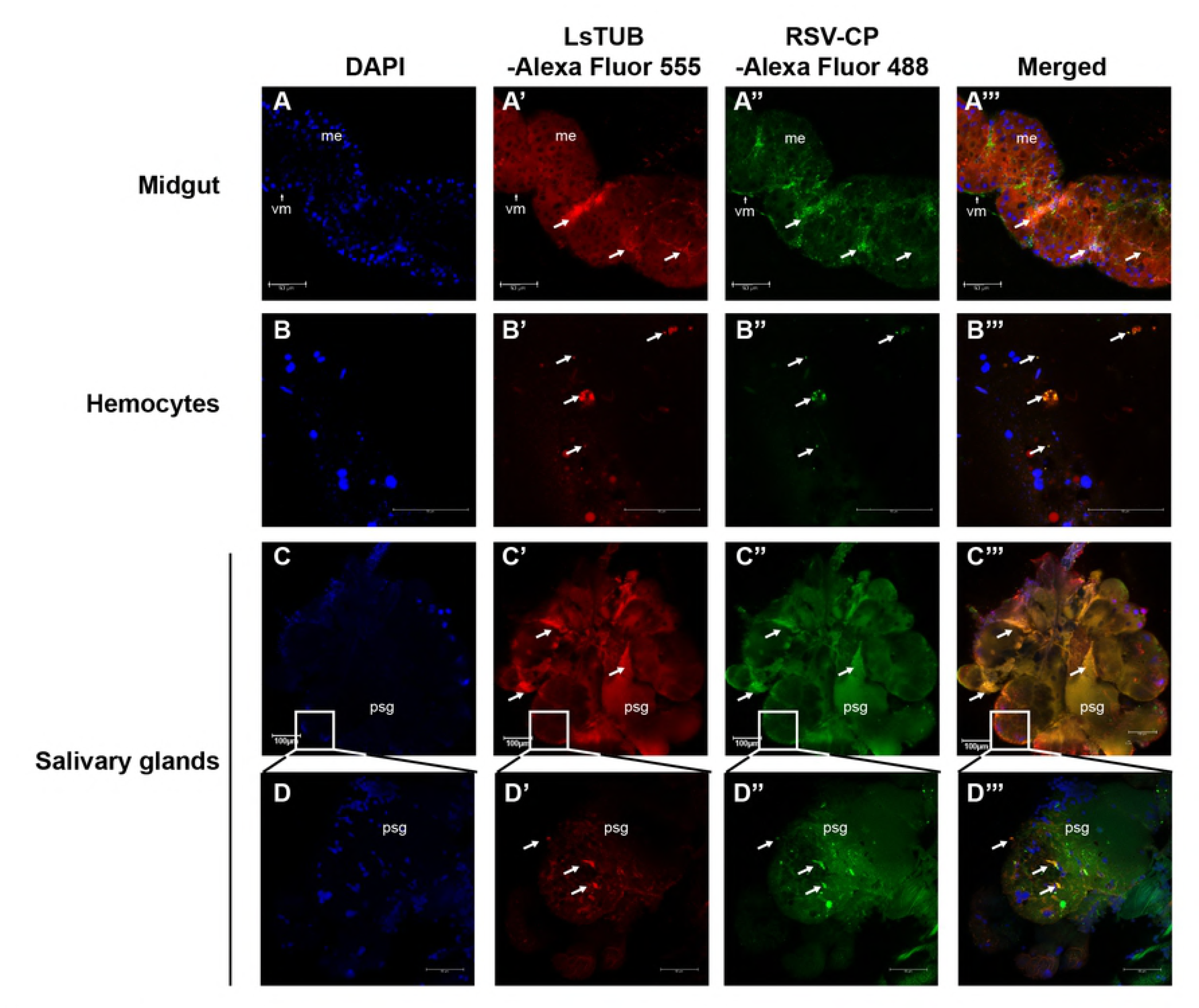
Localization of LsTUB and RSV in different tissues of SBPH. Detection of LsTUB and RSV coat protein (CP) in (A) SBPH midgut, (B) hemocytes, and (C) and salivary glands. Anti-LsTUB and anti-RSV CP monoclonal antibodies were conjugated to Alexa Fluor 555 (red) and 488 (green), respectively. Panels A-B’’’ and D-D’’’, bar=50 μm; panels, C-C’’’, bar=100 μm. Abbreviations: sg, salivary glands; psg, principal salivary glands; vm, visceral muscle; and me, midgut epithelium.

### LsTUB interacts with RSV NS3 in vitro

LsTUB was used as bait and a RSV cDNA library as prey in a yeast-two-hybrid (Y2H) assay designed to identify RSV proteins that potentially interact with LsTUB. The RSV nonstructural protein, NS3, interacted with LsTUB in the Y2H assay (Fig. 4A). To further examine the interaction between RSV NS3 and LsTUB, a pull-down assay was performed with glutathione S-transferase-tagged LsTUB (GST-TUB). When extracts from viruliferous SBPH were incubated with GST-TUB, western blot analysis indicated that NS3 co-immunoprecipitated with GST-TUB (Fig. 4B).

**Figure 4.**
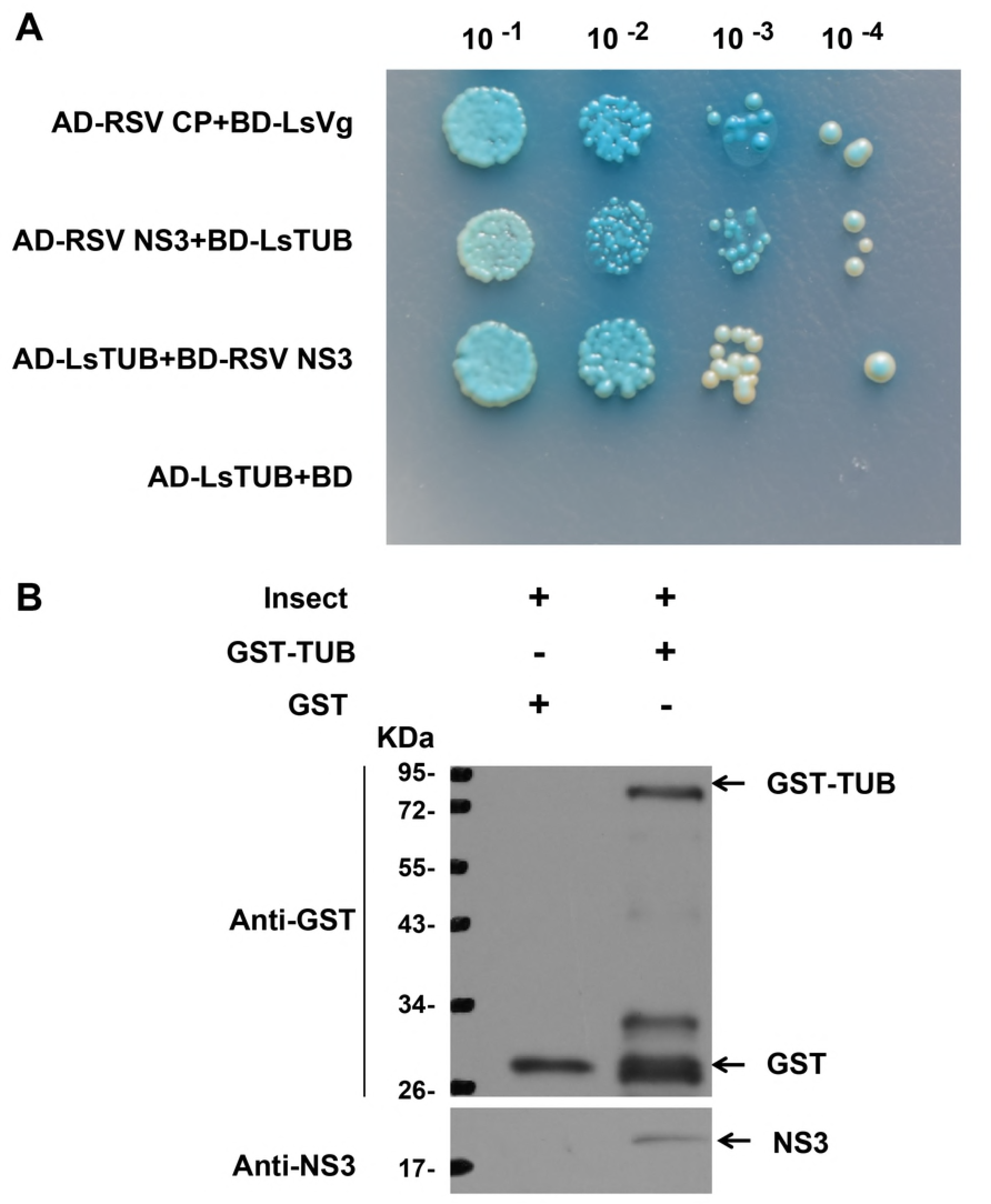
The interaction between LsTUB and RSV NS3 detected by yeast two-hybrid assay and GST pull-down assay. (A) Interactions between *L. striatellus* (LsVg and LsTUB) and RSV (CP and NS3) proteins using yeast two-hybrid assays. Yeast strain Y2HGold was cotransformed with RSV CP + LsVg (positive control) or LsTUB + NS3. Yeast cells were diluted 10^-1^ to 10^-4^ and plated onto QDO (SD-trp-leu-his-ade-20 mM3-AT) medium. Colonies growing on DDO were also assayed for β-galactosidase activity (blue color). AD-LsTUB + BD served as the negative control. Abbreviations: AD, activation domain, cloned in pGADT7; BD, bait domain, cloned in pGBKT7; CP, coat protein; LsVg, *L. striatellus* vitellogenin; LsTUB, *L. striatellus*, tubulin; NS3, nonstructural protein 3. (B) GST pull-down assays. LsTUB was fused to GST and incubated with viruliferous SBPH extracts or GST (control). Blots were probed with anti-NS3 or anti-GST antisera.

Considering that other RSV proteins may also bind LsTUB, we evaluated whether LsTUB could interact with RSV proteins: a putative membrane glycoprotein (NSvc2), capsid protein (CP), nonstructural disease-specific protein (SP) and movement protein (NSvc4), using Y2H analysis. Yeast strains containing full-length LsTUB as bait and the four proteins as prey failed to grow on synthetic dextrose dropout medium (data not shown). This result suggests that the interaction between LsTUB and NS3 was specific.

### Repression of *LsTUB* via RNAi reduces NS3 protein level and RSV titer in vivo

To further explore the potential role of LsTUB in NS3-mediated transmission of RSV, 3^rd^ instar viruliferous SBPH nymphs were supplied with dsRNAs derived from *GFP* (dsGFP) or *LsTUB* (dsTUB) via membrane feeding. After seven days of feeding, qRT-PCR showed that *LsTUB* mRNA in dsTUB-treated SBPHs was significantly reduced by more than 75% compared with the controls (untreated and dsGFP-treated SBPHs) (Fig. S1). These results indicated that RNAi-mediated knockdown of *LsTUB* was highly effective.

The midgut and salivary glands of viruliferous SBPHs were also examined by immunoblotting and confocal microscopy. Immunoblotting showed that dsTUB led to a decrease in LsTUB in both midgut and salivary glands, and this was also accompanied by a decrease in RSV-NS3 (Fig. 5A, B). Immunofluorescence and qRT-PCR indicated that dsTUB treatment caused a substantial reduction in RSV accumulation in the midgut and salivary glands. Furthermore, LsTUB and NS3 co-localized in tissues treated with dsGFP (see arrows), but not in tissues treated with dsTUB (Fig 5C-F’’’). Taken together, these results indicate that the interaction of LsTUB and NS3 is essential for RSV invasion and accumulation of the virus in the insect vector.

**Figure 5.**
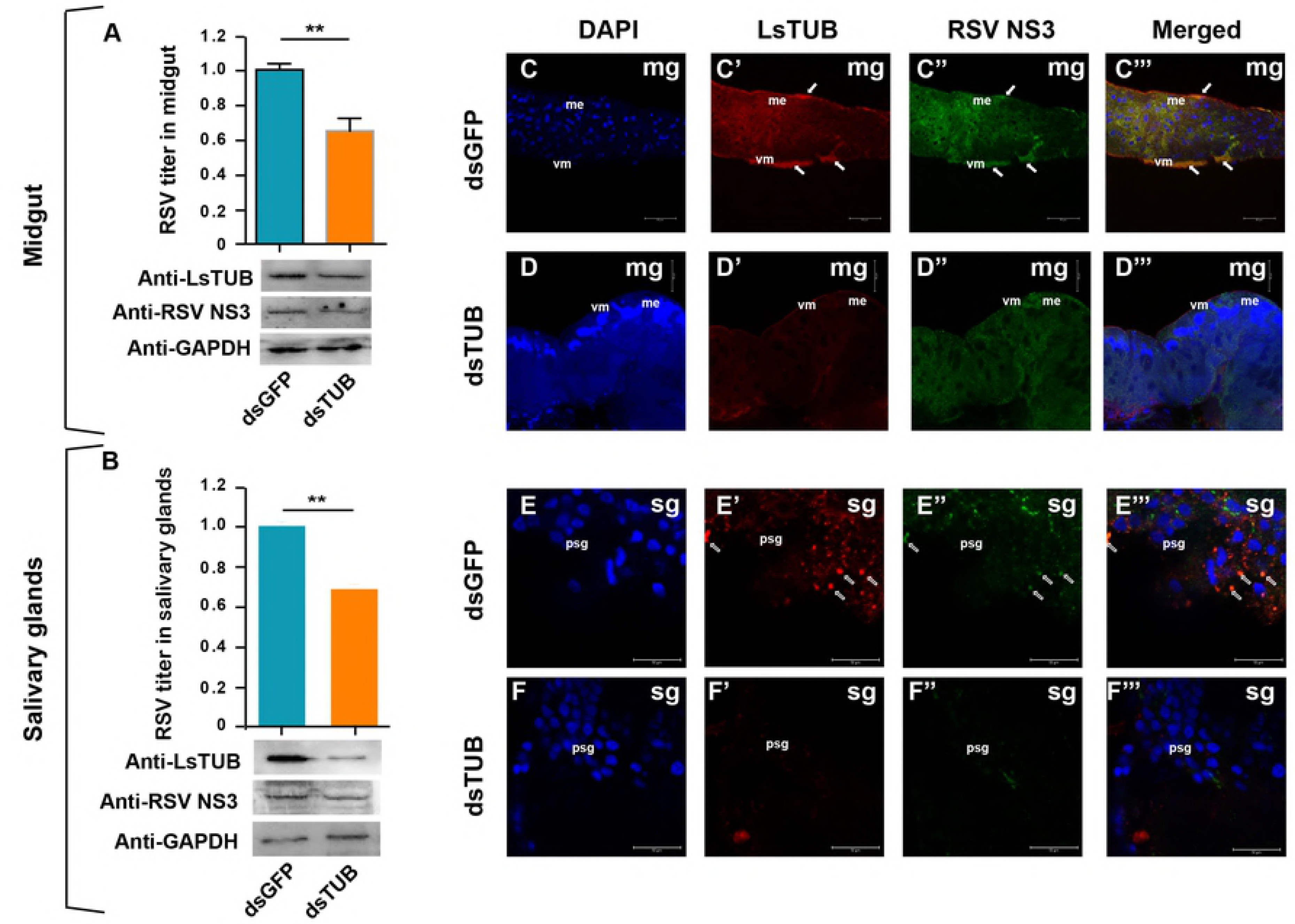
dsTUB-mediated RNAi inhibits RSV titers and reduces NS3 protein levels. Viruliferous SBPH were fed on artificial diets supplemented with dsGFP or dsTUB. SBPH midgut (A) and salivary glands (B) were analyzed for virus titer by qRT-PCR (Red columns indicates dsTUB; blue columns indicates dsGFP), and levels of LsTUB, NS3, and GAPDH were analyzed by immunoblotting. GAPDH was used as control. Each treatment was replicated three times for qRT-PCR, and values represent means ± S.E. A student’s T-test was used to analyze significance; ** represents *P*<0.01. The midgut (Fig. 5C-D’’’) and salivary glands (Fig. 5E-F’’’) of viruliferous SBPH were immunolabeled with anti-RSV CP (Alexa Fluor 488, green) and anti-LsTUB (Alexa Fluor 555, red) and then examined with confocal microscopy. Abbreviations: mg, midgut; sg, salivary glands; and vm, visceral muscle. Bar=50 μm.

### Knockdown of *LsTUB* results in decreased RSV transmission efficiency

The ability of dsTUB-treated viruliferous SBPH to transmit RSV was evaluated. Seven days after RNAi, viruliferous SBPHs treated with dsTUB or dsGFP were transferred to healthy rice seedlings, allowed to feed for two days, and then evaluated for virus inoculation rates by qRT-PCR (Table 1). At 15 days post-transmission, 12% of rice plants fed on by dsTUB-treated SBPH contained RSV, compared to over 40% of plants fed on by dsGFP-treated or untreated (CK) SBPH. The significance of transmission efficiency was evaluated by χ^2^ analysis, and *P*-values indicated that the inoculation rate of dsTUB-treated SBPH was significantly lower than the CK or dsGFP-treated viruliferous SBPH (Table 1). These results indicate that *LsTUB* plays a function in RSV transmission from SBPH to rice plants. Furthermore, RNAi-mediated knockdown of *LsTUB* inhibited horizontal transmission of the virus.

**Table 1.** Transmission rates of RSV-infected SBPH after RNAi.

| Treatment | Virus inoculation rates (%)<sup>a</sup> | $\chi^2$test<sup>c</sup> | | |
| --- | --- | --- | --- | --- |
| | | Treatments | $\chi^2$value | *P* value |
| Control | 0.40 | Cotrol vs dsTUB | 4.07 | 0.044 |
| dsGFP | 0.48 | dsTUB vs dsGFP | 4.67 | 0.031 |
| dsTUB | 0.12 | Control vs dsGFP | 0.29 | 0.588 |

a. The number of each treatment are the total plants inoculated from 20 independent RSV-infected insect.
c. There was a significant difference between the two treatments ( $P < 0.05$ ) by $\chi^2$ test.

### Effects of LsTUB knockdown on the feeding behavior of viruliferous *L. striatellus*

To investigate whether the decrease in RSV transmission efficiency was caused by RNAi-mediated changes in SBPH feeding behavior, the electrical penetration graph (EPG) technique was used to monitor SBPH feeding [28]. EPG signals were classified into seven different waveforms including NP, N1, N2-a, N2-b, N3, N4, and N5, which represent the following phases: non-penetration, penetration, stylet movement with salivary secretion, sustained salivary secretion, extracellular movement of the stylet around the phloem, phloem feeding, and xylem feeding, respectively [21]. Representative EPG waveforms after dsGFP and dsTUB treatment were not significantly different (Fig. 6), which demonstrates that LsTUB knockdown does not alter the feeding behavior of SBPH. Thus, the decrease in RSV transmission efficiency in dsTUB-treated viruliferous SBPH can be attributed to reduced RSV titers in the insect.

**Figure 6.**
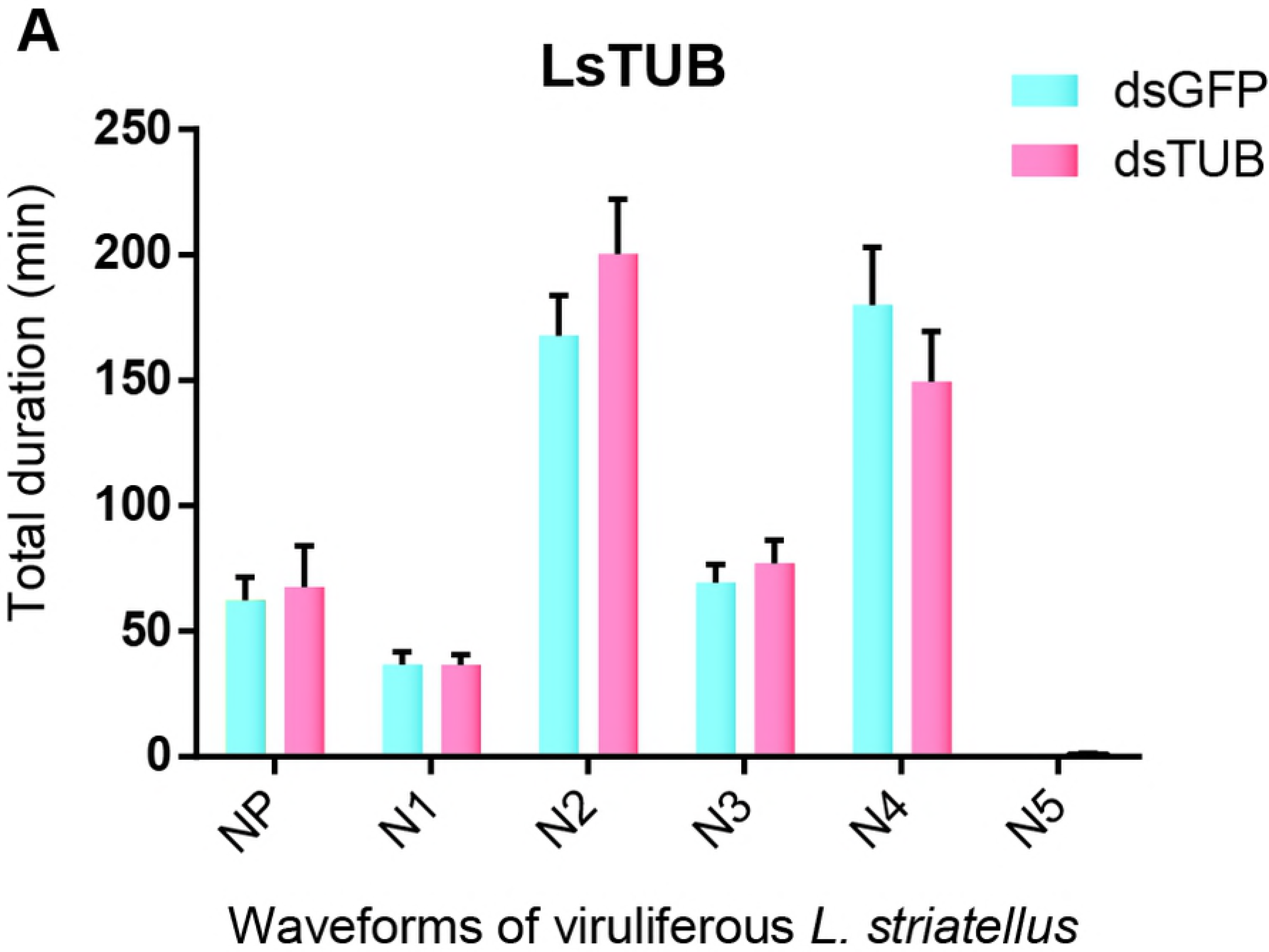
Electrical penetration graph (EPG) analysis of viruliferous *L. striatellus* fed on dsGFP or dsTUB. Waveform abbreviations: NP, non-penetration; N1, penetration; N2, salivation; N3, extracellular movement of stylet near the phloem; N4, sap ingestion in phloem; N5, water ingestion in xylem. Values represent means ± S.E; significance was evaluated by T-test analysis.

## Discussion

The intracellular protein tubulin is highly conserved in most organisms [29]. The tubulin family consisted of α, β and γ subfamilies, and the α- and β-tubulin subunits are highly heterogeneous and present numerous isotypes that vary in expression patterns [30]. Tubulin is part of the cytoskeleton and functions in intracellular transport and cell division in eukaryotic organisms [31]. Tubulin also functions in virus assembly and transport in various arthropods [25–27]. For example, the Dengue virus 2 envelope protein binds directly to tubulin or a tubulin-like protein in C6/36 mosquito cells [32]. β-Tubulin also serves as a receptor for swimming crab *Portunus trituberculatus* reovirus in crab [33]. Wen et al. [34] reported the existence of at least two α- and two β-tubulin genes in SBPH and obtained a full-length sequence of the SBPH gene encoding β-3 tubulin [34]. However, there are no prior reports documenting a role for SBPH tubulin in virus transmission. In our study, we cloned the gene encoding SBPH tubulin α-2 and show that it is highly expressed in RSV-infected salivary glands. Subsequent experiments showed that repression of *LsTUB* expression results in a decrease in RSV accumulation in midgut and salivary glands and inhibits RSV transmission to rice. These findings document a novel function for LsTUB in enabling RSV to overcome the midgut and salivary gland barriers of SBPH, which results in the dissemination of the virus to other organs in the insect vector.

RSV is an RNA virus containing four single-stranded genomes that use negative and ambisense strategies to encode the following proteins: RNA-dependent RNA polymerase, NS2, NSvc2, NS3, CP, nonstructural disease-specific protein (SP) and movement protein (NSvc4) [35–41]. Among these seven proteins, NS3 functions as a gene-silencing suppressor in plants and functions in the size-independent and noncooperative recognition of dsRNA [41,42]. A recent report demonstrated that RSV NS3 protein can hijack the 26S proteasome by interacting directly with the SBPH RPN3 protein [43]. Our observations using Y2H and GST pull-down assays confirmed that NS3 interacts with LsTUB (Fig. 4). Thus, addition to functioning as a gene-silencing suppressor, NS3 also interacts with LsTUB to enhance RSV dissemination in the insect vector.

The midgut and salivary glands of SBPH are important organs for RSV and represent the first and the last battlefields in the vector, respectively [17]. RSV particles initially establish infection in the midgut epithelium, then disseminate to the midgut visceral muscles, and ultimately move into the SBPH salivary glands where they can be introduced into rice. Whether RSV provokes different or similar reactions in the midgut and salivary gland of SBPH is a vital issue in understanding how the virus spreads in the insect vector and is re-inoculated into the host plant. Based on previous reports, two SBPH components are known to interact with the RSV CP to help the virus overcome the midgut barrier. One interaction consists of LsST6-CP binding, which mediates viral entry into midgut epithelial cells [18]. Another is GPS2-CP binding, an interaction that activates the SBPH JNK signaling pathway in the midgut, which is beneficial to viral replication [24]. With respect to the salivary gland barrier, only CPR1 or GPS2-CP binding were reported to facilitate viral movement in the salivary glands [23,24]. The spread of RSV in SBPH is obviously complex and requires multiple components. In this study, the interaction between SBPH tubulin and NS3 facilitated RSV accumulation and virus colonization of midgut and salivary glands, thus leading to successful inoculation of rice plants. Our findings thus complement and improve overall knowledge of the mechanistic basis of viral transmission in SBPH midgut and salivary glands.

Plant virus transmission is closely associated with the feeding behavior of insect vectors; therefore, monitoring the feeding process of dsRNA-treated SBPH can reveal the impact of dsRNA on feeding behavior and subsequent transmission to rice. Electrical penetration graph recordings have been used to investigate stylet penetration behavior in hemipteran insects [28,44]. In our study, the inoculation rate of dsTUB-treated SBPH was significantly lower than the control group; however, the feeding behaviors of the dsTUB-treated and control SBPH were not significantly different. Thus, the decrease in RSV transmission rate was not the result of altered feeding behavior, but is instead attributed to the failure of RSV to cross midgut and salivary gland barriers due to dsTUB treatment.

In summary, LsTUB helps RSV overcome the midgut and salivary gland barriers and enhances horizontal transmission of the virus. This conclusion is supported by immunofluorescent monitoring of LsTUB and RSV in midgut and salivary glands and by Y2H and pull-down assays with LsTUB and NS3 in vitro. Repression of LsTUB expression due to RNAi also reduced NS3 levels and consequently reduced viral dissemination into midgut and salivary glands, which ultimately reduced re-inoculation into the plant. These insights provide a better understanding of the interaction between plant viruses and vectors and may ultimately reveal new avenues for therapeutic intervention.

## Materials and methods

### Insects

RSV-free and viruliferous strains of *L. striatellus* were originally collected from Jiangsu province, China, and maintained in the laboratory for seven years. Both RSV-free and viruliferous *L. striatellus* were reared independently on seedlings of rice cv. Wuyujing 3 in glass beakers containing soil at a depth of 1 cm. Plants were maintained in a growth incubator at 25 ± 1°C, with 80% ± 5% RH and a 12-h light-dark photoperiod.

To ensure that insects were viruliferous, individual female insects were allowed to feed independently, and the resulting offspring were collected and analyzed via Dot-ELISA with the monoclonal anti-CP antibody [45]. Highly viruliferous colonies were retained and used in subsequent studies.

### Cloning and structure analysis of LsTUB

Approximately 100 salivary glands were dissected from *L. striatellus* adults and were considered as one sample; total RNA was isolated from the sample using TRIzol reagent and the manufacturer’s protocol (Invitrogen). The quality and concentration of total RNA were determined by spectrophotometry (NanoDrop, Thermo Scientific); 500 ng of RNA was subsequently used for reverse transcription in a 10 μl reaction with the PrimeScript™ RT reagent kit and gDNA Eraser as recommended by the manufacturer (Takara, Dalian, China). Based on the α-tubulin mRNA sequence downloaded from NCBI (GenBank accession no. AY550922.1), *LsTUB* cDNA was obtained using 5’- and 3’-RACE (Takara). The predicted LsTUB protein sequence was subjected to Blast analysis using DNAman software (LynnonBiosoft, USA), and domains of the predicted protein were deduced using SMART (http://smart.embl-heidelberg.de/) [46].

### RNA interference (RNAi)

The coding sequences of *LsTUB* and green fluorescent protein (GFP) were cloned into pMD19-T vectors (Takara, Japan). The primers for dsGFP and dsTUB amplification are listed in Table S1. Using the cDNA templates obtained above, dsRNAs were synthesized using the T7 RiboMAX™ Express RNAi System kit as recommended by the manufacturer (Promega, USA). A membrane feeding approach was used to introduce dsRNAs into SBPHs as described previously [47–49]. Briefly, second instar nymphs of viruliferous SBPH were maintained on a mixed diet containing 0.5 mg/ml dsRNAs for four days via membrane feeding and then transferred to healthy rice seedlings. The effects of dsRNA on *LsTUB* expression was evaluated by qRT-PCR.

### Real-time qRT-PCR

To measure *LsTUB* expression levels and RSV titers in SBPH, total RNA was isolated from whole SBPHs using the TRIzol Total RNA Isolation Kit (Takara, Dalian, China). Total RNA concentrations were quantified, and first-strand cDNA was synthesized as described above. The primers (Table S1) used for detecting RSV titers were designed based on *RSV CP*-specific nucleotide sequences. Similarly, *LsTUB* and *LsActin* (Control) primers (Table S1) was designed based on *LsTUB* and *LsActin* sequences. qRT-PCR was conducted using a CFX96™ Real-Time PCR Detection System (Bio-Rad, Hercules, CA, USA) and SYBR Premix Ex Taq (Takara, Dalian, China) as follows: denaturation for 3 min at 95 °C, followed by 40 cycles at 95 °C for 10 s, and 60 °C for 30 s. Relative expression levels for triplicate samples were calculated using the ΔΔCt method, and expression levels of target genes were normalized to the SBPH *Actin* gene.

### Western blotting

Whole body, midgut and salivary gland samples were collected and lysed to obtain total proteins. After adding 6× SDS loading buffer, samples were boiled for 10 min. The proteins were separated by 8-12% SDS-PAGE and transferred onto PVDF membranes. Blots were probed with anti-LsTUB antibody (1:1000 dilution), anti-RSV CP (1:1000 dilution), anti-RSV NS3 (1:500 dilution), or anti-GAPDH (1:2000 dilution). Immuno-reactive bands were detected using a goat anti-rabbit IgG-conjugated horseradish peroxidase (HRP) antibody and a goat anti-mouse IgG-conjugated HRP antibody (Proteintech, USA) at a 1:5000 dilution. Western blots were imaged with a Chemiluminescence Detection Kit (Bio-Rad, Hercules, CA, USA) and the Molecular Imager^®^ ChemiDoc™ XRS System (Bio-Rad).

### Immunofluorescence microscopy

SBPHs were maintained on rice plants for seven days after RNAi treatment. The salivary glands of SBPHs were dissected with forceps and fixed with 4% paraformaldehyde for 1 h. Samples were then blocked with fetal bovine serum (10%) at ambient temperature for 2 h. Preimmune serum and anti-LsTUB or anti-RSV CP were diluted 1:500 at 4 °C for 16 h and then visualized with Alexa Fluor 555- or Alexa Fluor 488-labeled secondary goat anti-rabbit IgG (CST, China). Salivary glands were then washed three times in PBS, and stained with 100 nM DAPI and CM-Dil (Sigma-Aldrich) for 2 min at room temperature. Fluorescence was observed with a Leica TCS SP8 STED confocal microscope (Leica, Germany).

### Yeast two-hybrid assay

Yeast two-hybrid assays were conducted using protocols supplied with the Yeastmaker™ Yeast Transformation System 2 (Takara-Clontech, USA). Briefly, the cDNA library of RSV was cloned as prey in pGADT7 using the Easy Clone cDNA library construction kit (Dualsystems Biotech), and full-length *LsTUB* was cloned as bait in pGBKT7. Positive clones were selected on SD quadruple-dropout (QDO) medium (SD/-Ade/-His/-Leu/-Trp), and interacting prey constructs were recovered and sequenced. To distinguish positive and false-positive interactions, we co-transformed empty pGADT7 and pGBKT7 into yeast strain Y2HGold, and ß-galactosidase activity was detected with the HTX Kit (Dualsystems Biotech).

### GST pull-down assay

*LsTUB* cDNA fragments were amplified and cloned into pGEX-3X as glutathione S-transferase (GST) translational fusions. Recombinant proteins were produced in *Escherichia coli* strain BL21 and purified. For pull-down assays, 1 mg viruliferous SBPH extract, 200 μl immobilized glutathione-Sepharose beads and 500 μg GST-LsTUB protein were added to 1 ml of pull-down buffer (50 mM Tris, 150 mM NaCl, 0.1% Triton X-100, 1 mM PMSF, 1% protease inhibitor cocktail [pH 8.0]), and then incubated at 4 °C for 16 h. Similarly, insect extracts were incubated with GST protein as a negative control. Beads were washed four times with pull-down buffer, and retained proteins were released by adding 2× loading buffer and incubating for 5 min at 95°C. Proteins were then separated by SDS–PAGE and detected with anti-GST (Cusabio, China) and anti-NS3.

### RSV transmission efficiency

SBPH adults derived from RSV-viruliferous strains were reared for five days on artificial liquid diets (50) amended with one of the following: dsGFP, dsLsTUB, or no additional substances (Control). Individual insects were cultured on healthy rice seedlings for two days, and the insect was then removed and analyzed by Dot-ELISA to detect whether it was viruliferous. If the insect was tested to be virus-free, the insect and corresponding seedling were both considered invalid for further study. The remaining rice seedlings infested with RSV-viruliferous insects were incubated another 10-15 days. And then whether the remaining rice seedlings was infected or not were determined by RSV CP primers (Table S1) via qRT-PCR. The number of viruliferous rice seedlings was recorded and calculated for transmission rates. Insects supplied with the dsGFP or CK diets were considered controls. Each viruliferous insect was considered to be one replicate and each treatment contained least 20 replicates. Transmission rates were calculated as transmission rate (%) = (number of infected seedlings/total number of seedlings) × 100. A χ^2^ test was performed with DPS statistical software (51) to detect differences between treatments.

### Electrical penetration graph (EPG) recording and data analysis

Nymphs from RSV-infected SBPHs fed on dsGFP or dsRNA were selected for this experiment. After a 30-min starvation, the mesonotum of *L. striatellus* was affixed with a gold wire (20 μm diameter, 2-3 cm long) using a soluble conductive adhesive. The *L. striatellus* individual was then connected to an eight-channel EPG recorder (Model: CR-8 DC-EPG I). Insects were placed on the culms of rice seedlings (three-leaf stage) in Ferrari insect cages; activity was recorded for 8 h in a greenhouse maintained at 25-26°C, with 60±5% RH. Insects were removed at the end of the 8-h period and analyzed for RSV by Dot-ELISA. If an insect tested virus-free, the data were considered invalid. Each treatment contained 20-30 replicates, and all recorded signals were analyzed.

## Figure Legends

**Figure S1. Relative levels of *LsTUB* mRNA after RNAi-mediated knockdown.**
*LsTUB* expression in untreated, dsGFP-, and dsTUB-treated SBPH. *LsTUB* expression was evaluated by qRT-PCR and normalized relative to GAPDH transcript levels. Values represent means ± S.E. Significance was evaluated by T-test analysis, and *** is significant at *P*<0.001. Treatments were replicated three times.

**Table S1.**
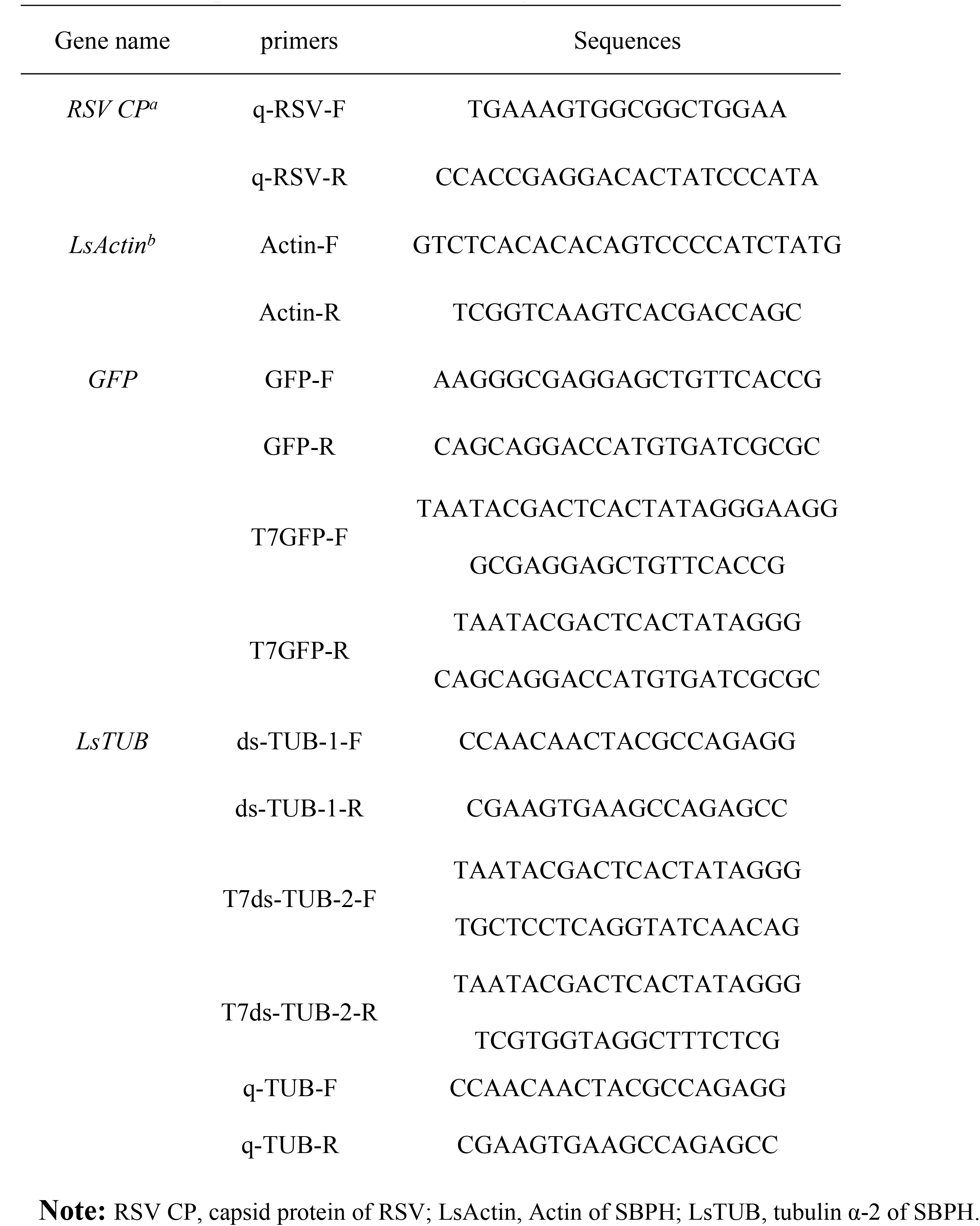
The primers used in this study.

